# Right Anterior Theta Connectivity Predicts Autistic Social Traits in Neurotypical Children

**DOI:** 10.1101/2022.03.26.485953

**Authors:** Aron T. Hill, Jodie Van Der Elst, Felicity J. Bigelow, Jarrad A. G. Lum, Peter G. Enticott

## Abstract

Growing evidence supports functional network alterations in autism spectrum disorder, however much less is known about the neural mechanisms underlying autistic traits in typically developing children. Using resting-state electroencephalographic (EEG) recordings, we examined whether functional connectivity could predict autistic trait expression in 127 children aged between 4-12 years. Regression models showed that right anterior theta connectivity was a significant predictor of autistic traits (p = 0.013), with increased connectivity in this region associated with greater autistic trait expression. These results corroborate similar recent findings in adults, extending this observation to a cohort of children spanning early-to-middle childhood. These findings further highlight EEG-derived functional connectivity as a sensitive physiological correlate of autistic trait expression in neurotypical children.

## Introduction

Efficient communication across disparate brain regions is crucial for cognition. This is made possible by synchronised activity across large populations of neurons, which can be recorded using electroencephalography (EEG) [1, 2]. Converging evidence from EEG studies indicates aberrant functional connectivity in autism spectrum disorder (ASD), providing a putative neurophysiological mechanism for the array of social and cognitive difficulties which often accompany this disorder [3]. Despite growing recognition of atypical patterns of connectivity in ASD, relative to neurotypical controls, it remains unclear if alterations in functional connectivity can predict autistic traits within the broader sub-clinical population. Autistic behavioural traits occur on a spectrum that extends beyond individuals with a clinical diagnosis and into the general population [4, 5]. That is, they are present within the wider population but at a level lower than is diagnosable as ASD, referred to as the broad autism phenotype [6]. In a recent study, Aykan et al. [7] provided compelling initial evidence of an association between EEG-derived resting-state theta connectivity and autistic traits measured across two separate groups of healthy adult participants. Specifically, these authors found that connectivity, as measured using magnitude-squared coherence, across electrodes spanning right anterior regions could predict autistic traits, measured using the Autism Spectrum Quotient (AQ), with higher connectivity values corresponding to greater autistic traits.

Resting-state EEG represents a convenient and cost-effective method for capturing spontaneous neural activity that is particularly amenable to research in children, who may be unable to perform more complex task-based paradigms [2]. The recent findings by Aykan et al. [7] compliment growing research implicating aberrant intrinsic neural activity patterns ASD [2, 3, 8]. The aim of the current study was to assess whether we could extend these findings to a population of typically developing children. To achieve this, we examined if connectivity could predict autistic traits measured using the Social Responsiveness Scale, 2^nd^ Edition (SRS-2), which is frequently used in research examining the broad autism phenotype, and shows convergent validity with the AQ [9, 10]. Additionally, to help protect against any potential confounds related to volume conduction of the neural signal, we utilised a robust analysis approach utilising weighted phase-lag index [wPLI] functional connectivity combined with spatial band-pass filtering (surface Laplacian) [11, 12]. In line with the results of Aykan et al. [7], we hypothesised that higher theta connectivity over the right anterior region would be associated with a greater degree of autistic traits.

## Methods

### Procedure

Data were collected during a single experimental session conducted either at the university laboratory, or in a quiet room at the participants’ school. Prior to commencement of the study, written consent was obtained from the parent or legal guardian of each child. Details of the experimental protocol were also explained to each child who then agreed to participate. Data reported in this study were collected as part of a larger neurocognitive and electrophysiological investigation into the development of the social brain in early and middle childhood [13].

### Participants

A total of 153 typically developing children, as described by their primary caregiver, were initially recruited to the study. Of these participants, 127 had complete EEG recordings and SRS-2 assessments and were included in the present analyses (see Table 1 for participant demographics). Ethical approval was provided by the Deakin University Human Research Ethics Committee (2017-065), while approval to approach public schools was granted by the Victorian Department of Education and Training (2017_003429).

**Table 1:** Demographic information and SRS scores

| Variable | Descriptive Statistics |
| --- | --- |
| N | 127 |
| Age (Mean ± SD) | 9.40 ± 1.94 |
| Gender (M/F) | 67/60 |
| WASI FSIQ (Mean ± SD) | 111.51 ± 11.95 |
| SRS Scores (Mean ± SD) | |
| SRS Total | 50.00 ± 9.95 |
| Social Awareness | 50.51 ± 9.88 |
| Social Cognition | 49.55 ± 10.28 |
| Social Communication | 50.01 ± 10.13 |
| Social Motivation | 50.13 ± 9.30 |
| Restrict. Int. Repet. behaviour | 49.71 ± 10.08 |

**Table 2:**
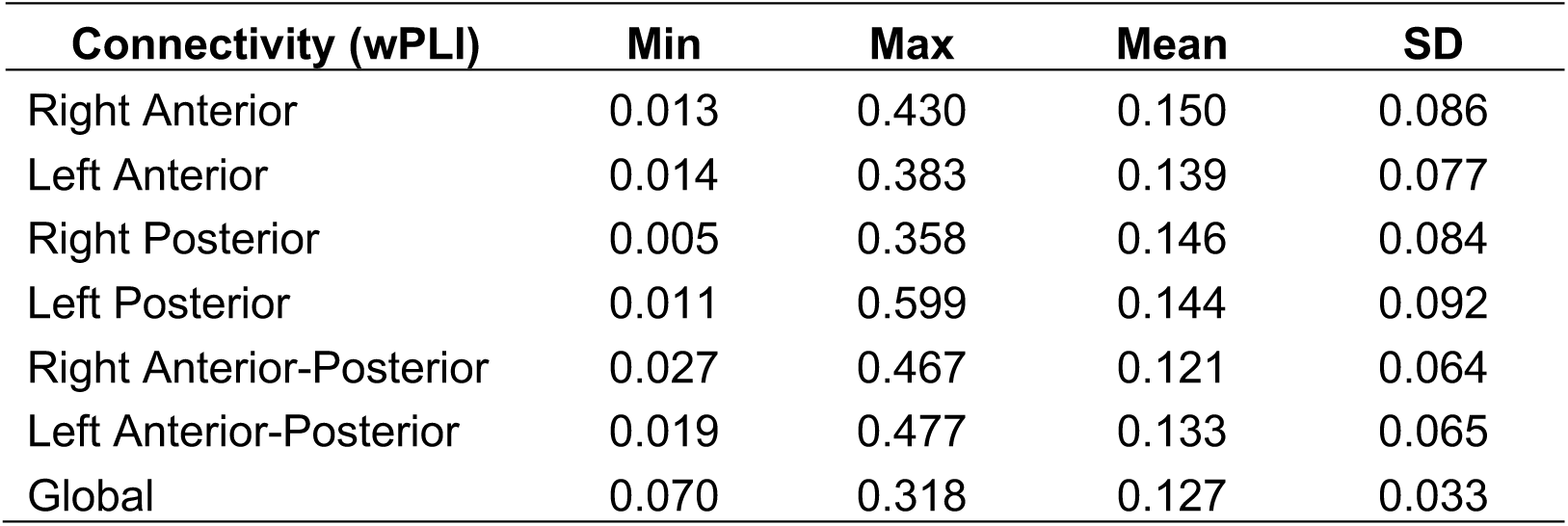
Theta connectivity values for each of the ROIs.

| Connectivity (wPLI) | Min | Max | Mean | SD |
| --- | --- | --- | --- | --- |
| Right Anterior | 0.013 | 0.430 | 0.150 | 0.086 |
| Left Anterior | 0.014 | 0.383 | 0.139 | 0.077 |
| Right Posterior | 0.005 | 0.358 | 0.146 | 0.084 |
| Left Posterior | 0.011 | 0.599 | 0.144 | 0.092 |
| Right Anterior-Posterior | 0.027 | 0.467 | 0.121 | 0.064 |
| Left Anterior-Posterior | 0.019 | 0.477 | 0.133 | 0.065 |
| Global | 0.070 | 0.318 | 0.127 | 0.033 |

### SRS-2

The SRS-2 is a 65-item rating scale that measures deficits in social behaviour associated with ASD [14, 15]. In the present study, the School-Age Form was used, which covers ages 4-18 years. Results are reported as T-Scores (Mean = 50, SD = 10), with higher scores indicative of greater autistic social impairment. The SRS-2 is comprised of a total score, which is considered the most reliable measure for social deficits associated with ASD [15], as well as five subscales tapping various dimensions of ASD-related traits (Social Awareness, Social Cognition, Social Communication, Social Motivation, and Restricted Interests and Repetitive Behaviour).

### EEG Recording and Analysis

Resting-state EEG data were recorded in a dimly lit room using a 64-channel HydroCel Geodesic Sensor Net (MagstimEGI, USA) containing Ag/AgCl electrodes surrounded by electrolyte-wetted sponges. Data were acquired using NetStation software (version 5.0) via a Net Amps 400 amplifier using a sampling rate of 1 KHz, with data online referenced to the Cz electrode. Prior to the commencement of recording, electrode impedances were checked to ensure they were < 50 KOhms. Data were recorded for two minutes while participants sat with their eyes open, and two minutes while participants had their eyes closed. In the present study, only the eyes-closed data were analysed so as to reduce the occurrence of electrooculography artifacts in the EEG record, as well as to remain consistent with Aykan et al. [7] who also reported results obtained from eyes-closed resting-state data. The EEG data were pre-processed in MATLAB (R2020a, The Mathworks, Massachusetts, USA) using an automated cleaning pipeline (Reduction of Electrophysiological Artifacts [RELAX]) [16]. For further details on data pre-processing see [17].

### Functional Connectivity

The pre-processed EEG data were segmented into three-second non-overlapping epochs. The estimate of the scalp current density (Surface Laplacian) was then obtained using the spherical spline method [18]. The EEG signal was then frequency transformed using a single Hanning taper to return the complex Fourier spectra for each subject/electrode for the theta band (4-8 Hz) and connectivity analysis was conducted using the weighted phase-lag index (wPLI) in Fieldtrip [19]. By using both a spatial filter (surface Laplacian) and the wPLI connectivity estimate, which disregards instantaneous (i.e., zero phase-lag) connections [12], we were able to provide a reliable estimate of phase synchronization between electrodes, and greatly reduce the likelihood of any spurious results caused by volume conduction [20, 21]. Consistent with Aykan et al. [7], functional connectivity was analysed across seven key regions: right anterior (F4-T8, C4-T8), left anterior (F3-T7, C3-T7), right posterior (C4-P8, P4-T8), left posterior (C3-P7, P3-T7), right frontal-parietal (F4-P4, F4-P8, F8-P4, F8-O8), left frontal-parietal (F3-P3, F3-P7, F7-P3, F7-P7). For each of these seven regions of interest, connectivity values across the electrode pairs were averaged, resulting in a single value per region for each participant which was used for statistical analyses. In addition, a ‘global’ connectivity estimate, representing the average connectivity value across all electrode combinations within the EEG cap, was also obtained for each participant.

### Statistical Analysis

All statistical analyses were performed in R (version 4.0.3; [22]). In keeping with the approach taken by Aykan et al. [7], we ran a stepwise linear regression to determine which of the connectivity values across the seven candidate regions (right anterior, left anterior, right posterior, left posterior, right fronto-parietal, left fronto-parietal, global) could predict autistic traits, as measured with the SRS-2 total score. Predictor variables were entered into the model if they demonstrated a p-value <.05, and were removed if they had a p-value >.10 using the ‘ols_step_both_p’ function from the ‘olsrr’ package. Residual diagnostics were also performed to assess assumptions for the regression model; this included visual inspection of residual Q-Q plots, residual versus fitted values plots, and histograms, as well as Shapiro-Wilk normality tests. As these revealed substantial deviations from normality (positive skew), the outcome variable was transformed using a Yeo-Johnson power transformation [23] prior to performing the analysis. Where the regression revealed an association between connectivity in any of the candidate regions and SRS total score, we then also ran additional Spearman correlations comparing connectivity values from that specific region and each of the SRS subscales. Multiple comparisons were controlled using false discovery rate (FDR) corrections [24].

## Results

Welch two-sample t-tests confirmed that there was no difference between males and females in terms of SRS total scores, t(116.26) = 0.849, p = 0.398, or intellectual function as measured using the Full Scale IQ from the Wechsler Abbreviated Scale of Intelligence, Second Edition (conducted in participants aged ≥6 years), t(119.38) = -0.741, p = 0.460. There were also no differences between males and females in terms of connectivity across any of the regions of interest, nor were there any significant associations between age and connectivity across any of the regions after FDR correction for the seven comparisons (all p_corrected_ > .05).

Connectivity values across the seven regions were then entered into a stepwise linear regression to determine the best predictor of SRS total score. This revealed a significant regression coefficient for the right anterior region, F(1,125) = 6.425, p = 0.013, adjusted R^2^ = 0.041, standardised β = 0.221. Given this significant finding, we then ran additional follow-up Spearman correlations comparing connectivity over the right anterior region to each of the five SRS subscales. This did not reveal any significant correlations after multiple comparison corrections across the five subscales (all FDR-corrected p > .05). However, trend-level associations between right anterior connectivity and social motivation (rho = 0.190, p_corrected_ = 0.080) and restricted interest and repetitive behaviours (rho = 0.218, p_corrected_ = 0.069) were observed (Figure 1).

**Figure 1:**
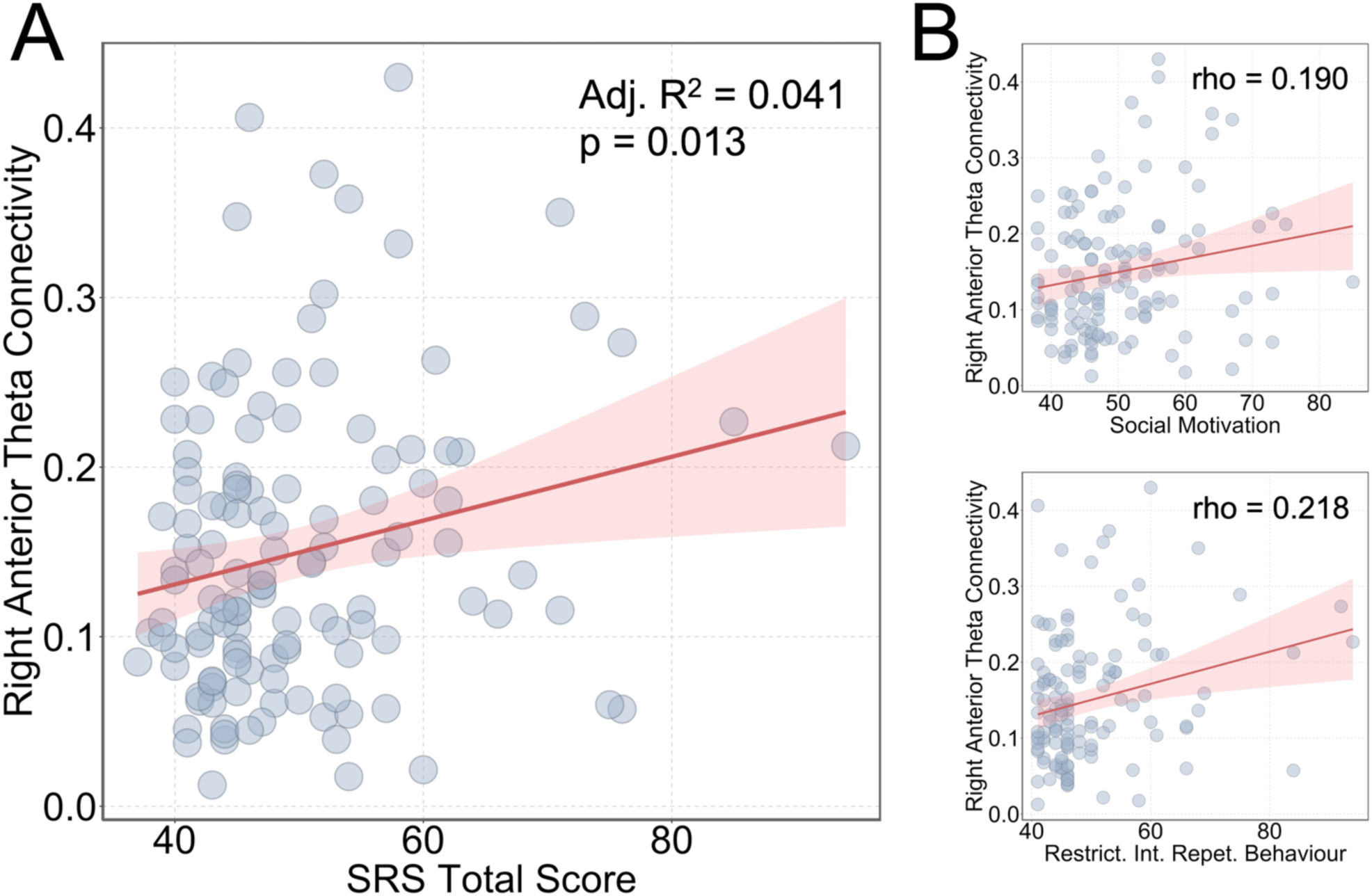
A) Scatterplot of theta connectivity over the right anterior region in relation to autistic social traits (SRS total score). B) Right anterior theta connectivity in relation to the SRS subscales: social motivation (top) and restricted interests and repetitive behaviour (bottom).

## Discussion

We assessed if functional connectivity within the theta band, extracted from resting-state EEG, could predict autistic social traits in a population of neurotypical children. We established a positive association between SRS-2 total scores and right anterior connectivity, indicating that increased connectivity was associated with more pronounced autistic trait expression. Further, by implementing a robust approach that utilised both a spatial EEG filter (surface Laplacian) as well as wPLI-based connectivity analysis, we were able to provide results that guarded against the influence of volume conduction effects, and thus are likely to provide a reliable non-invasive estimate of synchronized neural activity across the cortex [1, 12, 21].

The present results extend recent findings by Aykan et al. [7], who also demonstrated a link between right anterior theta connectivity and autistic traits, but in neurotypical adults. Despite a growing body of research indicating altered EEG connectivity patterns in ASD [8, 25], far less is currently known about how brain connectivity patterns change across the autistic dimension in sub-clinical populations. Our results provide initial evidence supporting a link between autistic traits and EEG-based resting-state measures of intrinsic functional connectivity in a population of children spanning early-to-middle childhood. This suggests that theta connectivity could play an important role in the processing of social information. The extant literature characterising functional connectivity patterns in ASD remains highly heterogeneous, with reports of both under- and over-connectivity across multiple brain regions [for review see: 3, 26]. Nevertheless, reduced neural synchronisation patterns have been observed in several previous studies comparing children with clinically diagnosed ASD to neurotypical controls [27-29]. The contrast between these previous observations, and our present findings, along with those of Aykan et al. [7], highlights potential differences between connectivity patterns across clinical and subclinical populations. However, we caution that additional research is needed to address this in more detail.

One potential explanation for our observed results, which was also suggested by Aykan et al. [7], is that increased theta connectivity might serve as a compensatory mechanism for individuals with high autistic traits, acting as a form of functional compensation for atypical development in neural circuitry within the brain. Whilst remaining speculative in relation to autistic traits, this concept has been raised by others [30], while more broadly, evidence for compensation via hyperconnectivity has been documented in patients following brain injury [31], and has been theorised to reflect a circuit-level response to neurological disruption [32].

Interestingly, despite connectivity being a significant predictor of SRS-2 total score, no significant associations were observed between connectivity and any of the five SRS-2 subscales after multiple comparison correction, suggesting that the severity of individual dimensions of autistic traits might not be robustly linked to connectivity. Nevertheless, the trend-level associations between connectivity and Social Motivation and Restricted Interests and Repetitive Behaviour (p = .08 and p = .069, respectively), warrant further investigation in future studies.

There were several limitations to the present study. First, while resting-state EEG recordings provide important insight into spontaneous neural activity within the brain, additional research is needed to explore associations between connectivity and autistic traits using-task based paradigms. Second, based on the observations of Aykan et al. [7], and in order to limit multiple comparisons, we restricted our connectivity analysis to the theta band. However, the broad range of EEG-related abnormalities reported in ASD [2, 3], suggest that additional work exploring potential associations between connectivity and autistic traits across wider frequency ranges is also warranted. Finally, although we utilised robust methods to reduce the influence of volume conduction effects on our results, future studies might further benefit from more advanced source-reconstruction techniques, which could also have the benefit of locating connectivity patterns at the cortical level [1, 33].

In conclusion, we have shown increased right anterior theta connectivity to be a neural correlate of autistic traits in neurotypical children. This extends recent research also linking right anterior theta connectivity to autistic traits in adults and highlights the utility of resting-state EEG for assessing functional connectivity in relation to autistic social traits in sub-clinical populations.

## Acknowledgement

The authors would like to thank Dr Gillian Clark for providing helpful feedback on this manuscript.

